# Repetition accelerates neural markers of memory consolidation

**DOI:** 10.1101/2022.12.14.520481

**Authors:** Wangjing Yu, Asieh Zadbood, Avi J. H. Chanales, Lila Davachi

## Abstract

No sooner is an experience over than its neural memory representation begins to be strengthened and transformed through the process of memory replay. Using fMRI, we examined how memory strength manipulated through repetition during encoding modulates post-encoding replay in humans. Results revealed that repetition did not increase replay frequency in the hippocampus. However, replay in cortical regions and hippocampal-cortical coordinated replay were significantly enhanced for repeated events, suggesting that repetition accelerates the consolidation process. Interestingly, we found that replay frequency in both hippocampus and cortex modulated behavioral success on an immediate associative recognition test for the weakly encoded information, indicating a significant role for post-encoding replay in rescuing once-presented events. Together, our findings highlight the relationships of replay to stabilizing weak memories and accelerating cortical consolidation for strong memories.

## Introduction

While many cognitive functions are under our conscious control, transforming everyday experiences into long-term memory is not one of them. Thus, how and when our experiences become stabilized in memory has been an intense topic of research. Ample evidence shows that the nature of these experiences—for example, whether they are emotional or involve reward motivation—modulates the probability of memories being retained after a delay (Phelps, 2004; Shohamy & Adcock, 2010; McGaugh, 2013). Further, the mechanisms underlying the benefits of emotional and reward-motivated memory are dependent on post-encoding consolidation processes (Phelps, 2004; Shohamy & Adcock, 2010; McGaugh, 2013). In contrast, relatively little is known about the dynamics and characteristics of memory consolidation processes for neutral experiences. Broadly speaking, existing work on memory consolidation shows that the neural activity associated with an experience is replayed during offline states after learning, both during quiet wakefulness and sleep (Skaggs & McNaughton, 1996; Foster & Wilson, 2006; O’Neill et al., 2010; Carr et al., 2011; Girardeau & Zugaro, 2011; Tambini & Davachi, 2013; Schapiro et al., 2018; Schuck & Niv, 2019; Liu et al., 2021). Importantly, post-encoding replay is closely associated with the ultimate fate of a memory (Girardeau et al., 2009; Staresina et al., 2013; Tambini & Davachi, 2013; Schlichting & Preston, 2014; Ólafsdóttir et al., 2018; Schapiro et al., 2018; Buch et al., 2021).

Another important characteristic of long-term memories is that, with consolidation, their representations become more distributed across the hippocampal-cortical networks (McClelland et al., 1995; Vilberg & Davachi, 2013; Squire et al., 2015). How memories become distributed remains unknown, but hippocampal-cortical replay is thought to be a leading observable mechanism that contributes to this process (Qin et al., 1997; Hoffman & McNaughton, 2002; Frankland & Bontempi, 2005; Ji & Wilson, 2007; Higgins et al., 2021; Tomé et al., 2022). Indeed, accumulating evidence in humans has shown that functional connectivity between the hippocampus and the cortex increases after novel learning and predicts memory retrieval success (Tambini et al., 2010; Tompary et al., 2015; de Voogd et al., 2016; Schlichting & Preston, 2016; Murty et al., 2017; Tambini & Davachi, 2019). A critical missing piece of the puzzle, however, is that we lack a comprehensive understanding of which factors modulate *offline replay dynamics* between the hippocampus and the cortex.

One way to approach this question is to examine the prioritization of recent experiences in post-encoding replay. While a consensus is emerging that memory consolidation is a highly selective process (Tompary et al., 2015; Gruber et al., 2016; Murty et al., 2017; Cowan et al., 2021), which *neutral* experiences the brain prioritizes for post-encoding replay remains largely unknown. On the one hand, from a Hebbian perspective, experiences associated with greater neural activity and co-activity during encoding (i.e. stronger encoding) should be more likely to replay or reactivate during post-encoding rest periods (Hebb, 1949; O’Neill et al., 2010). Outside of the domain of emotion and value, some prior work suggests that ‘strong’ memories are prioritized in replay. For example, a study by Tambini and Davachi (2013) decomposed aspects of the task signal during learning and showed that the strongest components of encoding activity patterns observed in the hippocampus selectively persist into the subsequent post-encoding rest period. On the other hand, some recent work in the context of a category learning paradigm demonstrated that weak, rather than strong, memories are prioritized during hippocampal post-encoding replay (Schapiro et al., 2018). Thus, taken together, there is mixed evidence regarding whether weakly encoded or strongly encoded memories are prioritized for replay.

Furthermore, in contrast to the growing literature on how hippocampal replay varies as a function of memory strength, little work has directly compared replay for strong and weak memories in the cortex. If memory distribution across hippocampal-cortical networks is a hallmark of memory stability and consolidation (McClelland et al., 1995; Vilberg & Davachi, 2013; Squire et al., 2015), it is also important to ask what features of new learning would modulate not only the frequency of replay events within the hippocampus and within cortical regions, but also the magnitude of coordinated replay *across* hippocampus and cortex.

Foundational behavioral work in humans has shown that one of the ways in which a neutral memory can be robustly strengthened is through repeated learning; decades of research has demonstrated that the frequency of encoding repetition is closely associated with later memory strength (Krueger, 1929; Wollen, 1962; Hintzman, 1970, 1976). However, the critical relationship between repeated encoding and post-encoding replay in the hippocampal-cortical systems is unknown. To address these open questions, in the current human fMRI study, we directly manipulated memory strength through repetition during encoding and then examined post-encoding replay of repeated (strong) and once-presented novel (weak) memories. To fully characterize differences in systems-level memory replay between conditions, we measured local hippocampal and cortical replay as well as coordinated hippocampal-cortical replay. Specifically, participants studied events that were either presented once (weak memory condition) or three times (strong memory condition) during encoding and this was followed by a post-encoding rest period. They were then tested on their memory for the studied pairs immediately following the rest period. Given that the encoding content was all visual in nature, we focused analyses on cortical replay in the higher-level visual processing region: ventral temporal cortex (VTC). We also selected the midline default network regions: retrosplenial cortex (RSC) and medial prefrontal cortex (mPFC), which are closely interconnected with the hippocampus (Mesulam, 1998; Vann et al., 2009; Ritchey & Cooper, 2020) and have been implicated in memory consolidation (Sterpenich et al., 2009; Nieuwenhuis & Takashima, 2011; Bonnici et al., 2012; Tompary & Davachi, 2017; Kaefer et al., 2022).

Our results show that repetition does not simply increase replay frequency in a non-specific manner. Instead, we find that post-encoding replay in cortical regions is significantly enhanced for repeatedly encoded information, while the hippocampus does not show differential replay across strong and weak memory conditions. Moreover, replay of strong memories was significantly more likely to be coordinated across hippocampal-cortical networks as compared to weak memories. This enhanced coordinated replay for strong memories suggests that repetition facilitates hippocampal-cortical communication and potentially accelerates consolidation processes across distributed brain regions. Interestingly, however, we also show that post-encoding replay explains behavioral variance in memory outcomes specifically for the weakly encoded information, indicating that while prioritizing strong memories, post-encoding replay compensates for weaker encoding and rescues weak memories.

## Results

### Cued-recognition memory performance

As expected, examination of cued recognition showed a significant memory benefit for thrice-presented over once-presented events. On average, participants selected the correct object given the word cue for an accuracy of 0.90 (SD=0.14) in the repetition condition and 0.69 (SD=0.20) in the once-presented condition. Both were significantly above chance (0.25; strong: t(28)=24.22, p<0.001, 95% CI[0.84, 0.95], Cohen’s d=4.50; weak: t(28)=11.98, p<0.001, 95% CI[0.62, 0.77], Cohen’s d=2.23; Figure 1C) and participants had significantly higher accuracy for the thrice-presented pairs as compared to the once-presented pairs (t(28)=5.82, p<0.001, 95% CI[0.13, 0.28], Cohen’s d=1.08; Figure 1C), confirming that repeated encoding improved memory performance (see Supplemental Information for results of response times on correct trials and for supplemental analysis comparing accuracy across encoding conditions and image categories).

**Figure 1.**
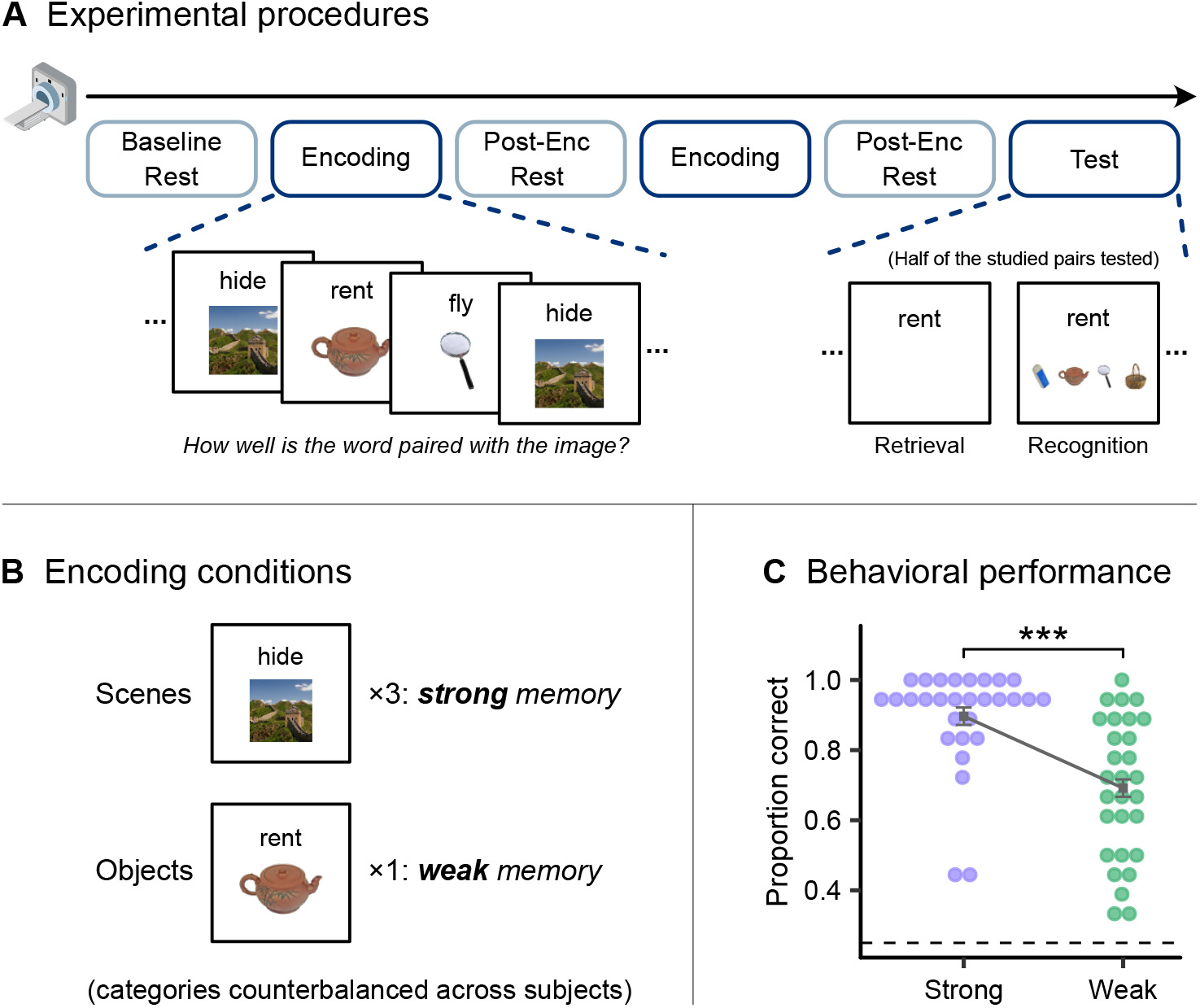
Experimental design and behavioral performance. (A) Experimental Procedures. The fMRI study started with a baseline rest period followed by the associative encoding of word-image pairs across two separate encoding blocks, each followed by a post-encoding rest period. Immediately after the second rest period, memory was tested on an associative recognition test for pairs from one of the encoding blocks. (B) During encoding, word-image pairs from one visual category (counterbalanced across participants) were presented three times, which formed the strong memory condition, and word-image pairs with images from another visual category were presented only once, which formed the weak memory condition. (C) Memory performance on the immediate associative recognition test. Each dot represents a participant, squared dots indicate mean accuracy. The dashed line represents the chance level (0.25). Error bars show within-subject standard error. ***p < 0.001.

### Post-encoding replay in the cortex prioritizes strong memory

Having confirmed that our manipulation led to a reliable difference in memory performance, we next measured post-encoding neural replay, or reactivation, for the strongly and weakly encoded memories. Specifically, for each subject and within each ROI (Figure 2A), we derived the multivoxel activity pattern associated with each encoding trial and with each time point (i.e., TR) in the subsequent post-encoding rest period. We then computed the neural pattern similarity (i.e., Pearson correlations; Kriegeskorte et al., 2008) between each encoding trial and all of the time points during the post-encoding rest periods. This resulted in an encoding-rest similarity matrix that represented the extent of pattern overlap between each encoding event and all time points during post-encoding rest periods (Figure 2B-C). Next, a threshold was applied to the matrix values in each rest time point to isolate encoding trials with the highest pattern similarity scores (i.e., exceeded 1.5 standard deviations above the mean of each given TR) as reliable evidence of neural ‘replay’ of these trials (Figure 2B-C; see Methods for details). This approach has been adopted in previous papers (Schapiro et al., 2018; Staresina et al., 2013) and was used as our primary approach to identifying neural replay. Note that a supplemental thresholding approach replicated the replay results. In this additional approach, the replay threshold for each encoding trial was derived from the pattern similarity distribution between each trial and all time points in the baseline rest period *prior to* encoding; we used the 95% percentile point (corresponding to a p-value of 0.05) in this ‘sham’ replay distribution as the post-encoding replay threshold for the trial (Staresina et al., 2013; see Methods for details).

**Figure 2.**
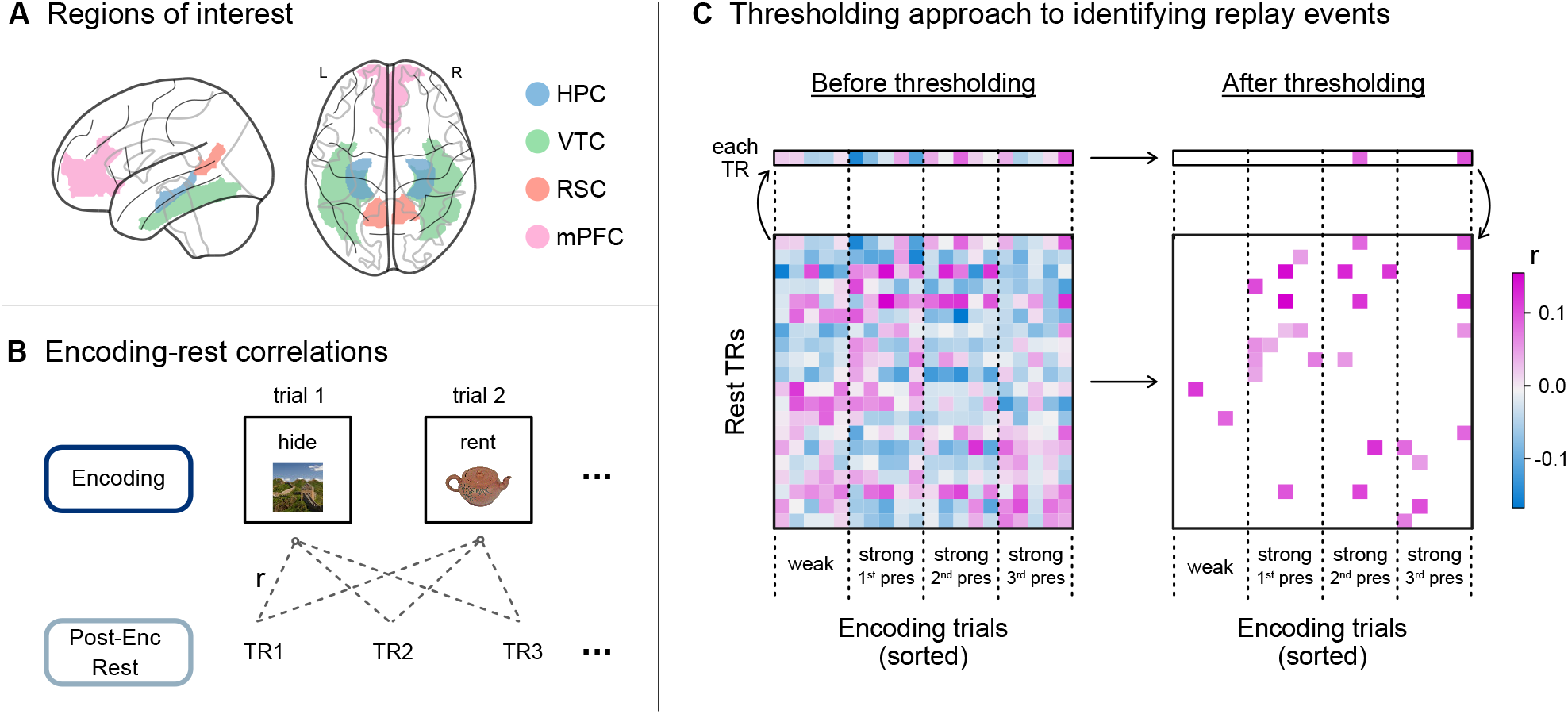
Replay analysis. (A) Regions of interest (ROIs) were defined within each participant. ROIs included the hippocampus (HPC) and ventral temporal cortex (VTC) extracted from Freesurfer’s volumetric segmentation (Fischl, 2012), as well as retrosplenial cortex (RSC) and medial prefrontal cortex (mPFC) converted from the Schaefer atlas (Schaefer et al., 2018). (B) To measure replay in each ROI, the activation pattern of each encoding trial was extracted and used to compute its similarity score (Pearson correlation coefficient) with the activation pattern of each timepoint, or TR, during the corresponding post-encoding rest. We then applied a thresholding approach as in panel C. (C) Schematic diagram of the thresholding approach using example matrices. We created a correlation matrix for each pair of encoding block and post-encoding rest period in each ROI and for each participant. We then thresholded the matrix TR by TR to select only similarity scores greater than 1.5 SD above the mean score as evidence for replay of the encoding trials (Staresina et al., 2013; Schapiro et al., 2018).

After obtaining the thresholded matrix (Figure 2C right), the number of rest time points that showed reliable replay evidence of encoding trials within the weak and strong conditions were used to calculate the replay frequency for each condition (see Figure S1 for supplemental analysis comparing replay across both encoding conditions and image categories). Importantly, for the strong memory condition, replay frequency was computed separately for each of the three presentations: strong 1^st^ presentation, strong 2^nd^ presentation, and strong last presentation. Our main comparison of interest was between the weak memory condition and strong *last presentation* condition (see reasons discussed below; see Methods for details).

Perhaps somewhat surprisingly, we found no evidence for significant differences in the rate of replay for strong and weak memories in the hippocampus (t(28)=-0.60, p=0.56, 95% CI[4.89, 2.69]; Figure 3, top panel, highlighted). Given that there is evidence for functional distinctions across the hippocampal long axis (Jung et al., 1994; Poppenk et al., 2013; Brunec et al., 2018; Thorp et al., 2022), we performed a follow-up analysis of replay within the anterior and the posterior thirds of the hippocampus. The results did not reveal any significant differences in replay frequency across conditions in the anterior (t(28)=-2.01, p=0.11, 95% CI[-7.71, 0.64]) or the posterior hippocampus (t (28)=1.41, p=0.34, 95% CI[-1.32, 5.19]).

Next, we examined post-encoding replay in select cortical regions. Replay frequency in these cortical regions was robustly and significantly enhanced for the repeated memories compared to once-presented events (Figure 3, bottom panel, highlighted). This was evident in all three cortical ROIs: the ventral temporal cortex (VTC: t (28)=5.73, p<0.001, 95% CI[10.74, 27.92], Cohen’s d=1.06), the retrosplenial cortex (RSC: t (28)=5.61, p<0.001, 95% CI[8.23, 21.91], Cohen’s d=1.04), and the medial prefrontal cortex (mPFC: t(28)=5.80, p<0.001, 95% CI[8.30, 21.29], Cohen’s d=1.08). These results provided strong evidence that repeated encoding increased the frequency of memory replay in cortical networks as compared to the once-encoded experiences (see replicated results with the baseline rest thresholding approach in Figure S2) and this was in contrast to the replay pattern in the hippocampus.

For the preceding analyses, we focused on comparing replay of once-presented items with the replay of the *last presentation* of the thrice-presented items. There were two reasons why we selected the strong memory *last presentation* condition. First, by selecting only one iteration of presentations within the strong memory condition, we ensure that the comparison of replay frequency between strong and weak memory conditions was not biased by the total number of encoding trials in each condition. Second, we reasoned that memory representations for repeatedly presented pairs are changing every time participants study these pairs during encoding, and that this change in representation with repetition may be reflected in post-encoding replay. Specifically, we hypothesized that for the repeatedly studied pairs, post-encoding replay may be more likely to prioritize the most recent, or the newest, presentation.

To directly examine this hypothesis, we next computed replay frequency across the three presentations (strong 1^st^, 2^nd^ and last presentations) of trials in strong memory condition, treating them as separate conditions. In all cortical ROIs, there was a significant difference in the replay frequency across the three repetitions of the strong memory conditions (VTC: F(2, 56)=25.69, p<0.001, η^2^=0.41; RSC: F(2, 56)=29.56, p<0.001, η^2^=0.27; mPFC: F(2, 56)=40.13, p<0.001, η^2^=0.43) (Figure 3, bottom panel). We showed that the last presentation of the thrice presented items was most frequently replayed across the three repetitions, followed by strong 2^nd^ presentation condition, while trials in the strong 1^st^ presentation condition were least replayed). These results provided strong evidence that the most updated, or newest representations, of repeatedly encountered information are more likely to be replayed in cortical regions. By contrast, this prioritization of memory for the last presentation was not evident in the whole hippocampus (F(2, 56)=1.32, p=0.28). Follow-up analyses of the hippocampal subregions showed that there was also no difference in the anterior third of the hippocampus (F(2, 56)=0.24, p=1), but a marginal difference was noted in the posterior hippocampus such that there was a trend towards a significant increase in replay for the newest representations of the repeated trials (F(2, 56)=3.76, p=0.059, η^2^=0.096; Figure 3, top panel).

Thus far, our results show clear evidence for increased post-encoding replay of strongly encoded memories compared to the weak ones in midline cortical regions and ventral temporal cortex. Further, replay events tended to reflect the most updated memory representation measured as the last presentation of the thrice presented items was most replayed. One concern with this latter result, however, is whether or not our measures of replay were merely driven by the temporal proximity between encoding and rest. In other words, on average the last presentation of the thrice-presented items occurred closer in time to the post-encoding rest period where we measured replay events, while the once-presented items often occurred evenly across the whole encoding blocks. To control for the temporal lag between item encoding and post-encoding rest, we performed a supplemental control analysis where we compared replay of a subset of trials from each of the strong (last presentation) and weak memory conditions that were matched in terms of their temporal proximity to the post-encoding rest period. For each participant, we selected the latter half of the weak memory trials in each encoding block; then for each selected weak memory trial, we identified strong memory (last presentation) trials within the same block that was presented closest in time and *prior to* the given weak memory trial, ensuring that overall the selected weak memory trials were in fact encoded *more recently* (i.e., closer in time to the post-encoding rest period) than the selected strong memory trials (see Methods for details).

**Figure 3.**
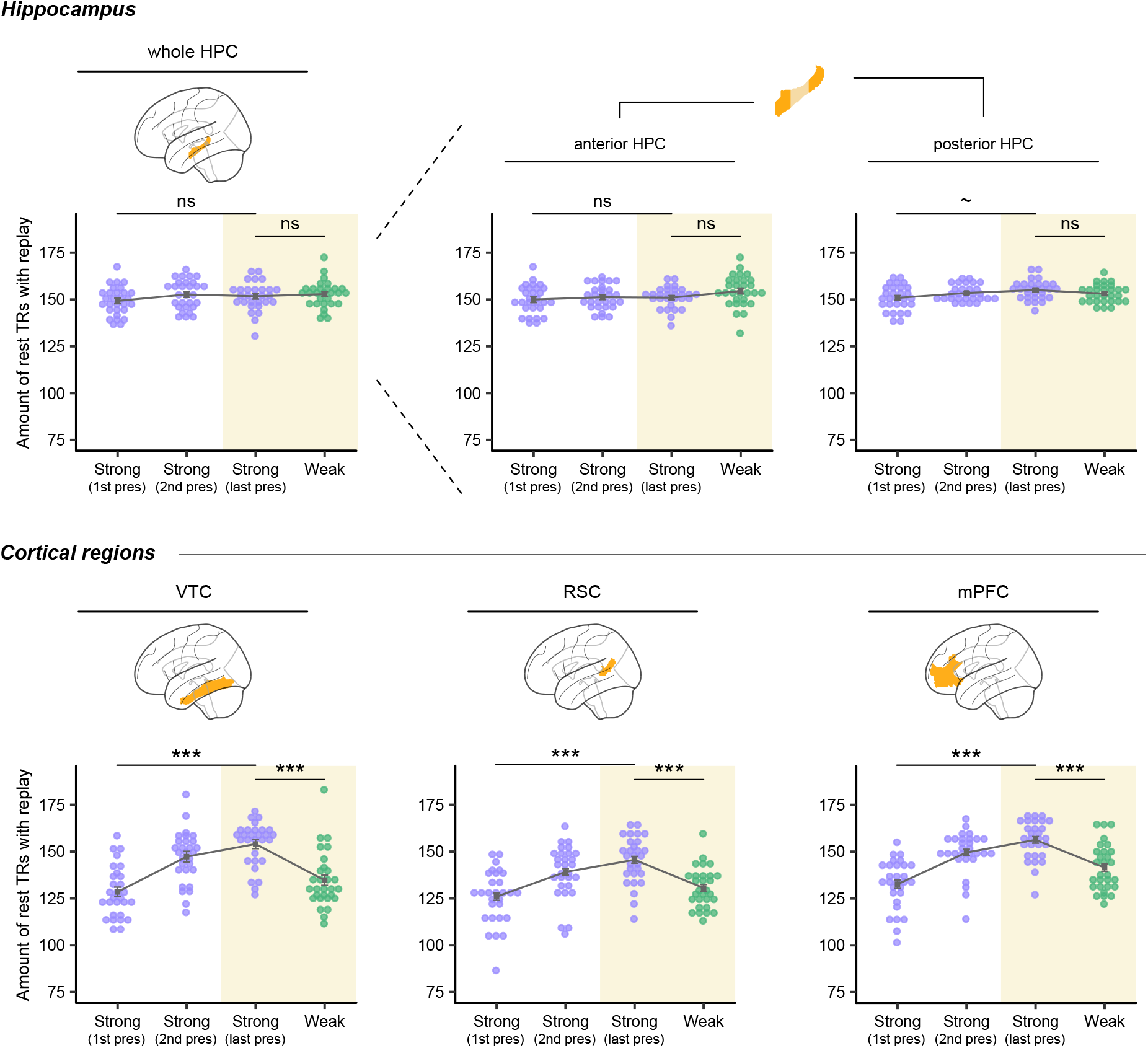
Post-encoding replay of strong and weak memories. Post-encoding memory replay for each of the three presentations of the repeated trials (strong memories) and the once-presented trials (weak memories), in the hippocampus (top panel) and cortical ROIs (bottom panel). The main comparison of interest was between strong last presentation condition and weak memory condition, which was highlighted in yellow. Each dot represents a participant, squared dots indicate mean values. Error bars show within-subject standard error. ns: not significant, ∼p<0.1, ***p < 0.001 (statistical significance adjusted with Bonferroni correction).

Results revealed that in all cortical ROIs, the rate of replay across strong (last presentation) and weak memory conditions persisted after controlling for encoding recency. This was seen significantly in the ventral temporal cortex (t (24)=2.27, p=0.032, 95% CI[0.90, 18.70], Cohen’s d=0.45) and the medial prefrontal cortex (t (24)=2.35, p=0.027, 95% CI[1.43, 21.77], Cohen’s d=0.47), and marginally in the retrosplenial cortex (t (24)=1.87, p=0.073, 95% CI[-0.84,17.24], Cohen’s d=0.37) (Figure S3). It is worth noting that a decrease in the effect sizes of these results was expected due to the reduction of power as fewer trials were included in this analysis (see Methods for detail). Nevertheless, these findings provide important evidence that the prioritization of strong memory in post-encoding replay in the cortex is a phenomenon not be fully explained by encoding recency.

### Coordinated hippocampal-cortical replay

After assessing replay in the hippocampus and cortical regions separately, we next examined whether replay across the hippocampus and the cortex were differentially synchronized for strong and weak memories. This was inspired by prior work showing that memories learned in a distributed fashion are associated with significantly enhanced hippocampal-cortical connectivity during their reactivation but not increased hippocampal or cortical activation locally (Vilberg & Davachi, 2013). Therefore, while the hippocampus did not prioritize strong over weak memories for replay, we hypothesized that repetition may accelerate coordinated replay in addition to local cortical replay frequency.

Extensive literature has suggested that different hippocampal subregions are not only functionally distinct from each other (Jung et al., 1994; Poppenk et al., 2013; Brunec et al., 2018; Thorp et al., 2022), but importantly, they also show differently weighted connectivity with cortical regions (Insausti & Muñoz, 2001; Kahn et al., 2008; Aggleton, 2012; Adnan et al., 2016; Ritchey & Cooper, 2020). Therefore, in examining coordinated replay with cortical regions, we separately examined each of the anterior and posterior thirds of the hippocampus. For each rest TR and within each memory condition, we took the binary replay patterns across the encoding trials (replayed, or not replayed) from the thresholded similarity matrix of each brain region (see Figure 2C right for an example matrix). We then computed the Jaccard similarity (Jaccard, 1912) between the patterns for each pair of hippocampal and cortical regions (e.g. anterior hippocampus and VTC) during that TR, which represented the proportion of trials that were replayed *simultaneously* in both regions. We then obtained an averaged Jaccard similarity index across all rest time points for each condition and for each participant, as a metric of coordinated replay (see Methods for details).

Results revealed significantly more coordinated memory replay between the posterior hippocampus and all cortical ROIs for strong memories than weak memories (Figure 4; posterior hippocampus & VTC: t (28)=3.57, p=0.0039, 95% CI[0.0025, 0.015], Cohen’s d=0.66; posterior hippocampus & RSC: t (28)=3.06, p=0.015, 95% CI[0.00094, 0.010], Cohen’s d=0.57; posterior hippocampus & mPFC: t (28)=3.28, p=0.0084, 95% CI[0.0012, 0.0098], Cohen’s d=0.61). These results demonstrate that repetition increases the proportion of coordinated replay events between the posterior hippocampus and cortex. Similar numerical patterns were also found between the anterior hippocampus and cortex, but the difference across conditions didn’t survive Bonferroni correction (all t(28)<2.19 and p>0.11). Additionally, for coordinated replay with ventral temporal cortex and with medial prefrontal cortex, the difference between strong and weak memories was significantly greater for posterior, as compared to, anterior hippocampus (VTC: F(1,28)=58.79, p<0.001, η^2^=0.32; mPFC: F(1,28)=24.96, p<0.001, η^2^=0.21). This interaction between hippocampal subregion and memory condition was not found for the coordinated replay with the retrosplenial cortex (F(1,28)=0.69, p=1). Taken together, these results show that hippocampal-cortical coordinated replay is enhanced for repeated memories. Importantly, this effect appeared to be regionally specific, such that it was statistically stronger between posterior, as compared to anterior, hippocampus and each of the ventral temporal cortex and medial prefrontal cortex, which was not the case for retrosplenial cortex.

**Figure 4.**
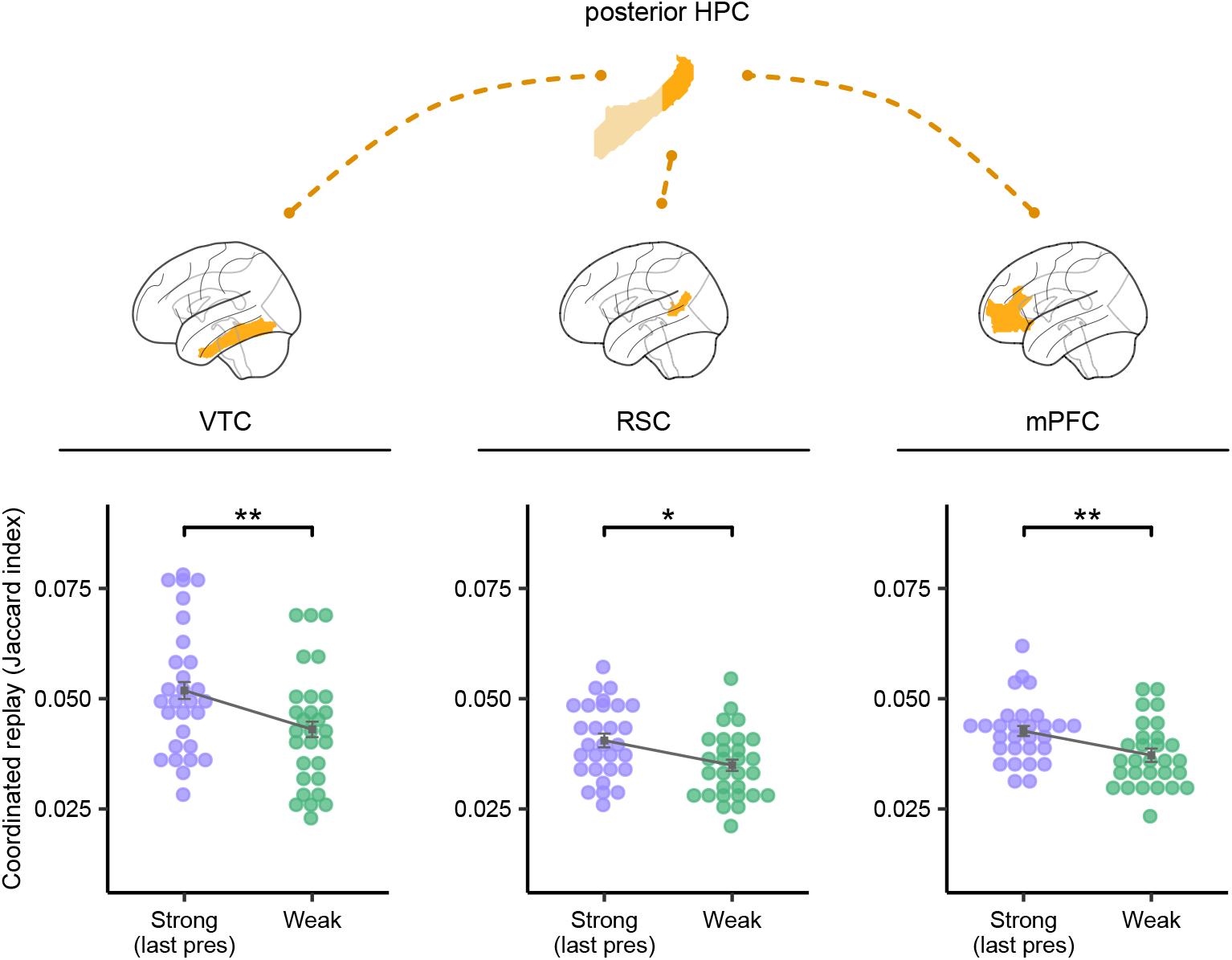
Coordinated replay between the posterior hippocampus and each of the cortical ROIs for strong and weak memories. We computed Jaccard similarity index between replay patterns across the posterior HPC and each of the cortical ROIs as our metric of coordinated replay, which represented the proportion of trials that are simultaneously replayed in both regions. Each dot represents a participant. Squared dots indicate mean values. Error bars show within-subject standard error. *p<0.05, **p < 0.01 (statistical significance adjusted with Bonferroni correction).

### Post-encoding memory replay and behavior

Post-encoding resting state activity has been suggested to support the consolidation of memories which has, in turn, been associated with their behavioral accessibility (Tambini et al., 2010; Deuker et al., 2013; Staresina et al., 2013; Tambini & Davachi, 2013, 2019; Schlichting & Preston, 2014; Tompary et al., 2015; Gruber et al., 2016; Murty et al., 2017; Poskanzer et al., 2021). Here, we asked if post-encoding neural replay was related to memory behavior for weakly and strongly encoded memories. To investigate this question, we ran a trial-level mixed-effects linear model in each ROI, predicting retrieval success of each tested pair (remembered or forgotten on the immediate memory test) with the replay frequency of the corresponding encoding trial in the strong (last presentation) or weak memory conditions during the post-encoding rest period, while also controlling for the trial-specific univariate activity during encoding (see Methods for details; see Supplemental Tables for full model outputs).

Results revealed a significantly positive association between trial-specific replay frequency in the hippocampus and retrieval success probability during the memory test selectively for weakly encoded pairs (b=0.044, p=0.036; Figure 5, top panel; Table S1-1). This result suggests that post-learning hippocampal replay explains variance in memory outcomes seen at the time of the memory test for once-presented, or weakly encoded, information. Breaking the whole hippocampus into anterior and posterior portions, we found a similar pattern in the anterior portion, which showed a significantly positive association between replay and memory specifically in the weak memory condition (b=0.056, p=0.0042; Figure 5, top panel; Table S1-1). By contrast, the association between replay in the posterior hippocampus and memory outcomes was not significant in either weak or strong conditions (both p>0.13; Figure 5, top panel; Table S1-1 & S1-2). Together, these results suggest hippocampal replay, in the anterior portion especially, contributes to the successful retention of weakly encoded information. In a follow-up exploratory analysis focused on individual differences (see Methods for details), we also found that post-encoding replay was positively associated with memory for the once presented trials in ‘poor performers’, but not in ‘good performers’ whose performance was closer to ceiling. This result potentially indicates that replay is especially essential for those who showed limited memory for information with one-shot encoding opportunity (Figure S4; Table S3).

Examination of cortical regions, by contrast, did not show strong evidence for an association between cortical replay in ventral temporal cortex or medial prefrontal cortex and memory outcomes (all p>0.19; Figure 5, bottom panel; Table S2-1 & S2-2). However, in the retrosplenial cortex, the association between replay frequency and memory for the once-encoded pairs was significantly modulated by the levels of univariate activity during the encoding of that pair (replay frequency × univariate activity: b=-0.00065, p=0.0091; Figure 5, bottom panel; Table S2-1). Specifically, replay frequency in the retrosplenial cortex more positively predicted retrieval success of the once-presented items associated with *weaker* univariate encoding activity. In line with our finding that hippocampal replay rescues once-presented memories, cortical replay in the retrosplenial cortex differentially rescues the weakest of the weak memories, that is, those once-presented items associated with low univariate activity during encoding. We did not find that replay in hippocampus or any cortical region significantly predicted immediate recognition success of the repeatedly encoded memories (all p > 0.13; Table S1-2 & S2-2). We proposed a few accounts that may explain this lack of association, which we addressed in the Discussion.

**Figure 5.**
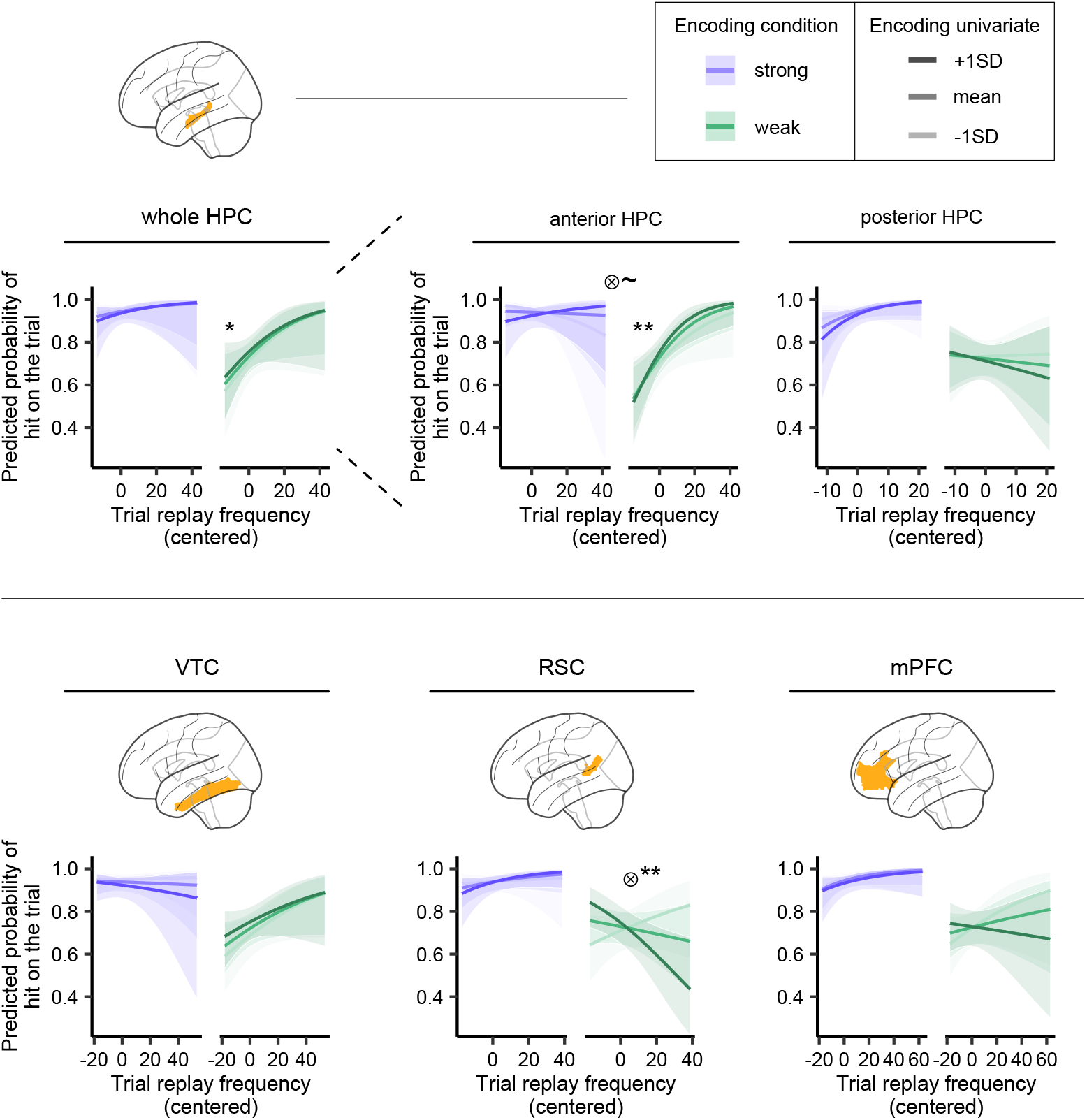
Predicting memory outcome with post-encoding replay. Mixed-effects linear models predicting memory outcome of each pair with the trial-specific replay frequency and univariate encoding activity in weak and strong (last presentation) memory conditions. Saturation of lines indicates the strength of encoding univariate activity (included as a continuous variable in the models). Ribbons represent 95% confidence intervals. ⊗ interaction, ∼p<0.1, *p<0.05, **p<0.01 (statistical significance adjusted with Bonferroni correction).

## Discussion

In the current human fMRI study, we examined how the frequency and nature of post-encoding memory replay are modulated by differential learning dynamics. Specifically, we directly manipulated memory strength during encoding by presenting study events either once or three times. We then computed the frequency of replay events in the hippocampus and cortical regions as well as how often those replay events were coordinated after this learning. Interestingly, our manipulation did not increase replay frequency in the hippocampus, which was statistically equivalent for once- and thrice-encoded memories. However, we found a striking increase in memory replay of repeated events in the cortical regions examined, which include the ventral temporal cortex and midline cortical regions: retrosplenial cortex and medial PFC. Moreover, repeated encoding also resulted in increased coordinated replay between hippocampus and cortex. These results indicate that repetition during learning accelerates systems-level memory consolidation by promoting memory distribution across the hippocampus and cortex, a known biomarker of memory stabilization (McClelland et al., 1995; Vilberg & Davachi, 2013; Squire et al., 2015). Importantly, our findings highlight that memory stabilization, as facilitated by memory replay, can be modulated by our decisions about how and when we study new materials.

While cortical replay and hippocampal-cortical coordinated replay were significantly increased for repeated compared to once-presented items, the frequency of replay events for once-presented items was related to behavioral measures of immediate memory. Specifically, we found that post-encoding replay in hippocampus was significantly correlated with immediate memory outcomes of once-presented events, but not for repeated events. Furthermore, replay in retrosplenial cortex was most protective for once-presented items with low levels of encoding activity. These findings reveal that while post-encoding replay reflects the strength of learning (repeated > once-encoded memories), at the same time, it may be most critical for rescuing the weakest memories (Schapiro et al., 2018). Consistent with this adaptive rescue perspective, our exploratory analyses reveal that across participants, post-encoding replay benefits weak memories specifically in ‘poor’ rather than ‘good’ performers. These results are also broadly in line with recent findings in rodents showing that offline replay predicts memory stability of spatial information more for reward-distant than reward-adjacent locations, suggesting that replay contributes to the consolidation even for information that is less central to the learning experience (Grosmark et al., 2021). Supporting this view, prior research on sleep and memory has also revealed that weakly encoded memories benefit more from sleep-based consolidation (Denis et al., 2021). Collectively, these findings indicate that offline processing helps maintain memories that could otherwise likely to be forgotten.

Some prior work has shown, using principal components analyses, that the strongest components of a learning event tend to be the ones that persist into post-encoding rest (Tambini & Davachi, 2013). Additionally, memories associated with high emotional valence (de Voogd et al., 2016), high reward (Gruber et al., 2016; Liu et al., 2021; Murty et al., 2017) or high utility (Terada et al., 2021) have been shown to be prioritized during memory consolidation. Consistent with these findings, we show that neutral memories strengthened through repeated encoding are also more extensively replayed in the cortex compared to events with more limited encoding opportunity. In everyday life, as compared to single occurrences, events that repeatedly appear are more likely to be of behavioral relevance. It is possible that repetition during learning increases plasticity in the associated neural ensembles that may serve as a ‘tag’ for replay.

Importantly, we also show that coordinated replay between the posterior hippocampus and the cortex is significantly enhanced for the repeatedly encoded memories as compared to the once-encoded memories. We hypothesize that the increased level of coordinated replay is associated with the distribution of memory across the hippocampal-cortical networks. This idea is consistent with the notion that memory representations become less hippocampal-dependent and more distributed throughout the cortex with consolidation (McClelland et al., 1995; Frankland & Bontempi, 2005; Takashima et al., 2009; Girardeau & Zugaro, 2011; Vilberg & Davachi, 2013; Squire et al., 2015; Robin & Moscovitch, 2017; Antony et al., 2017). Here, we propose that as a result of memory distribution, the hub of replay may also shift from the hippocampus to the cortex. Our findings suggest that this process can be accelerated by repeated encoding.

Perhaps surprisingly, in the current study, we do not find convincing evidence in the hippocampus for differential replay between strong and weak memories. By contrast, and as mentioned above, some prior work in humans has demonstrated that the hippocampus selectively replays the strongest components of the neural encoding activity patterns during post-encoding rest (Tambini & Davachi, 2013), while others have shown that hippocampal replay prioritizes information associated with worse memory performance (Schapiro et al., 2018). One possible reason for the different findings may be the inconsistent definition or manipulation of memory strength. In the current study, memory strength was directly manipulated during encoding via repeated presentations, while prior work has distinguished strong and weak memories with other criteria. Using this repetition manipulation, we see that the benefits of repeated encoded lie solely in increasing cortical replay and hippocampal-coordinated replay. This is consistent with the idea that repeated encoding accelerates the consolidation process. During post-encoding rest, memories of repeated events are not only more replayed in the cortical regions, but may also be farther along in the process of being distributed in the cortex. In contrast, memories for the once-encoded events are lagging behind in cortical replay and thus may still be highly hippocampal-dependent. Consistent with this possibility, we see that hippocampal replay specifically rescues memories for these once-presented items. Our results are also consistent with findings from a recent study in rodents, that reported active hippocampal replay following single experiences and decreased hippocampal replay rate for repeated experiences (Berners-Lee et al., 2022).

Critically, the repetition manipulation in our experimental design also allowed us to examine replay for separate presentations of each repeated event. This analysis showed that replay in the cortex prioritized the most updated, or the newest, memory representations, as compared to memory for the earlier presentations of the same content. This is an important finding as it highlights that our memory systems prioritize the replay of the most updated encounters, because this information may be more relevant for future goals (c.f., Gillespie et al., 2021). Existing work has demonstrated that when the same information is repeatedly studied, a new memory trace is established each time it is encoded; this was evidenced by people’s ability to discriminate between multiple occurrences of the same information (Hintzman & Block, 1971; Hintzman, 1988). Extending prior literature, the current study shows that different memory representations of a repeated experience are also associated with varying levels of post-encoding replay. These representations may be separate and distinct, and thus, memory for each presentation of a repeated event may also go through diverging consolidation processes.

Interestingly, in the current study, we did not find any significant association between replay frequency and immediate memory outcomes for the repeated events. This lack of association could be due to several reasons. First, in the current study, memory was tested immediately following the rest period and memory performance was close-to-ceiling levels in the strong memory condition in many participants (mean accuracy = 0.90). Therefore, the limited variance in the trial-to-trial performance in the strong memory condition may have prevented us from effectively capturing a relationship between post-encoding replay and retrieval success.

Thus, in order to see the benefits of immediate replay on later memory for repeated events, experiments would need to measure stability over time by the inclusion of both an immediate memory probe as well as one after a longer delay (Litman & Davachi, 2008; Vilberg & Davachi, 2013). Another possibility is that accelerated consolidation for repeatedly encoded memories, as evidenced by enhanced cortical and hippocampal-cortical replay, may also be leading to memory integration with prior knowledge (Schlichting & Preston, 2015; Tompary & Davachi, 2017), or memory transformation towards gist-like representations (Robin & Moscovitch, 2017; Antony et al., 2021). However, the associative recognition test we implemented here did not target behaviors associated with these processes. Future investigations are needed to examine precisely how replay shapes memory representations.

In conclusion, the current study provides important evidence that repeated encoding leads to significantly increased levels of cortical replay and hippocampal-cortical coordinated replay. This was not the case for hippocampal replay, which did not statistically differ for once- and repeatedly-encoded memories. These findings suggest that repetition accelerates memory consolidation by facilitating the distribution of memory across the hippocampal-cortical networks. We also reveal that post-encoding replay explains more variance in the subsequent behavioral outcomes of weakly encoded memory, demonstrating that replay adaptively contributes to the stabilization of weak memory. Together, our findings provide novel insights into the hippocampal-cortical dynamics during the process of memory stabilization. Future work can further investigate other aspects of our learning experience and assess how and when we study materials impact memory consolidation by examining post-encoding replay dynamics.

## Methods

### Participants

A total of 32 paid participants were recruited and completed the full experiment (mean age=23.0 years old; 24 females and 8 males). All participants were right-handed, reported normal or correct-to-normal vision, and provided written informed consent. Three participants were excluded from all analyses (two participants were excluded due to excessive head motion; one participant was excluded due to substantial signal dropouts in cortical regions). The final sample consisted of 29 participants (mean age = 23.1 years old; 23 females and 6 males). The study protocol was approved by the New York University Committee on Activities Involving Human Subjects.

### Materials and Procedures

The full experiment consisted of a total of three sessions spanning three days. Note that only the first session of the full experiment was within the scope of the current study. The first session consisted of a functional localizer block, two encoding blocks, a test block and a restudy block, together with a rest period at the beginning of the session (baseline rest) as well as immediately following each of the encoding, test, and restudy blocks, yielding a total of 5 rest periods. All parts of this session were conducted in an fMRI scanner. However, only the baseline rest period, the two encoding blocks, and their corresponding post-encoding rest periods, as well as the test block were relevant to the current study (Figure 1A). Therefore, details of the other blocks in the first session and the other two experimental sessions on day 2 and day 3 are not further reported in this paper.

The stimuli used in the current study included 144 words and 108 images. The words were commonly used verbs with a length between 3 and 11 letters (mean length = 5.71 letters). The images consisted of colored photographs of three visual categories: objects (e.g., teapot), faces (e.g., Beyoncé), and scenes (e.g., Forbidden City); there were 36 images in each visual category. Each of the 108 images was then paired with a word; the pairing between words and images was randomized across participants. Each participant studied word-image pairs with images from two of the three visual categories. Each of the two presented categories was assigned to strong or weak memory condition (see “Encoding”). The selection of the two presented visual categories (out of objects, faces, or scenes), as well as the assignments of categories to encoding conditions (strong, weak), were counterbalanced across participants.

#### Encoding

Each participant studied 72 words paired with images from two visual categories. The word-image pairs were equally split across two encoding blocks, which had an identical structure. Within each encoding block, pairs with images from one visual category were presented only once, which formed the weak memory condition; whereas pairs with images from the other category were presented three times, which formed the strong memory condition (Figure 1B). Each encoding block included a total of 72 trials (36 different pairs, 18 pairs in each memory condition), presented in a randomized order. Each encoding trial was on screen for 4s, during which participants were asked to rate how well the word was associated with the image using a button box. Each trial was then followed by a 6s intertrial interval (ITI), during which a fixation cross was presented.

#### Test

Participants completed an associative recognition test immediately following the encoding phase. They were tested on memory for pairs studied from one of the two encoding blocks (pairs from another encoding block were studied again in a restudy block, which was irrelevant to the current study). The temporal interval between the memory test and the corresponding encoding block was controlled for across all participants. The test included a total of 36 trials. Each test trial started with a 4s retrieval phase, in which a studied word was presented on screen and participants were asked to mentally recall the image paired with the word. Following the retrieval phase, participants were asked to make an associative recognition judgment about the pair. Four image choices appeared on the screen together with the word, and participants were asked to choose the image that was paired with the word by pressing the corresponding button using a button box. The word and the image choices were presented on screen for 2s, but the response window continued into a 6s ITI following each trial.

#### Rest periods

The study began with a 7-minute rest scan (baseline rest period; 210 TRs), followed by the first encoding block. Each of the two encoding blocks was also immediately followed by another 7-minute rest period (post-encoding rest period; 210 TRs each). Participants were asked to keep their eyes closed while remaining awake during all rest periods.

### MRI data acquisition

Imaging data were acquired on a 3T Siemens scanner with a Siemens head coil at the Center for Brain Imaging at New York University. Anatomical images were collected using a T1-weighted protocol (256 × 256 × 240 matrix; 0.9 mm^3^ voxels). Functional images were collected with a T2*-weighted gradient-echo EPI sequence (repetition time = 2s; echo time = 24.2ms; flip angle = 68°; 92 × 92 × 60 matrix; 2.25 mm^3^ voxels; multiband acceleration factor = 2). We also collected in-plane field map scans to improve co-registration between anatomical and functional images.

### fMRI data preprocessing

Results included in this manuscript come from preprocessing performed using fMRIPrep 1.5.9 (Esteban, Markiewicz, Blair, et al., 2018; Esteban, Markiewicz, Goncalves, et al., 2018; RRID:SCR_016216), which is based on Nipype 1.4.2 (Gorgolewski et al., 2011; Gorgolewski et al., 2018; RRID:SCR_002502).

#### Anatomical data preprocessing

The T1-weighted (T1w) image was corrected for intensity non-uniformity (INU) with N4BiasFieldCorrection (Tustison et al., 2010), distributed with ANTs 2.2.0 (Avants et al., 2008, RRID:SCR_004757), and used as T1w-reference throughout the workflow. The T1w-reference was then skull-stripped with a Nipype implementation of the antsBrainExtraction.sh workflow (from ANTs), using OASIS30ANTs as target template. Brain tissue segmentation of cerebrospinal fluid (CSF), white-matter (WM) and gray-matter (GM) was performed on the brain-extracted T1w using fast (FSL 5.0.9, RRID:SCR_002823, Zhang et al., 2001). Volume-based spatial normalization to two standard spaces (MNI152NLin2009cAsym, MNI152NLin6Asym) was performed through nonlinear registration with antsRegistration (ANTs 2.2.0), using brain-extracted versions of both T1w reference and the T1w template. The following templates were selected for spatial normalization: ICBM 152 Nonlinear Asymmetrical template version 2009c [Fonov et al., 2009, RRID:SCR_008796; TemplateFlow ID: MNI152NLin2009cAsym], FSL’s MNI ICBM 152 non-linear 6th Generation Asymmetric Average Brain Stereotaxic Registration Model [Evans et al., 2012, RRID:SCR_002823; TemplateFlow ID: MNI152NLin6Asym].

#### Functional data preprocessing

For each of the 20 BOLD runs found per subject (across all tasks and sessions), the following preprocessing was performed. First, a reference volume and its skull-stripped version were generated using a custom methodology of fMRIPrep. A B0-nonuniformity map (or fieldmap) was estimated based on two (or more) echo-planar imaging (EPI) references with opposing phase-encoding directions, with 3dQwarp Cox and Hyde (1997) (AFNI 20160207). Based on the estimated susceptibility distortion, a corrected EPI (echo-planar imaging) reference was calculated for a more accurate co-registration with the anatomical reference. The BOLD reference was then co-registered to the T1w reference using flirt (FSL 5.0.9, Jenkinson & Smith, 2001) with the boundary-based registration (Greve & Fischl, 2009) cost-function. Co-registration was configured with nine degrees of freedom to account for distortions remaining in the BOLD reference. Head-motion parameters with respect to the BOLD reference (transformation matrices, and six corresponding rotation and translation parameters) are estimated before any spatiotemporal filtering using mcflirt (FSL 5.0.9, Jenkinson et al., 2002). The BOLD time-series (including slice-timing correction when applied) were resampled onto their original, native space by applying a single, composite transform to correct for head-motion and susceptibility distortions. These resampled BOLD time-series will be referred to as preprocessed BOLD in original space, or just preprocessed BOLD. The BOLD time-series were resampled into several standard spaces, correspondingly generating the following spatially-normalized, preprocessed BOLD runs: MNI152NLin2009cAsym, MNI152NLin6Asym. First, a reference volume and its skull-stripped version were generated using a custom methodology of fMRIPrep. Several confounding time-series were calculated based on the preprocessed BOLD: framewise displacement (FD), DVARS and three region-wise global signals. FD and DVARS are calculated for each functional run, both using their implementations in Nipype (following the definitions by Power et al., 2014). The three global signals are extracted within the CSF, the WM, and the whole-brain masks. Additionally, a set of physiological regressors were extracted to allow for component-based noise correction (CompCor, Behzadi et al., 2007). Principal components are estimated after high-pass filtering the preprocessed BOLD time-series (using a discrete cosine filter with 128s cut-off) for the two CompCor variants: temporal (tCompCor) and anatomical (aCompCor). tCompCor components are then calculated from the top 5% variable voxels within a mask covering the subcortical regions. This subcortical mask is obtained by heavily eroding the brain mask, which ensures it does not include cortical GM regions. For aCompCor, components are calculated within the intersection of the aforementioned mask and the union of CSF and WM masks calculated in T1w space, after their projection to the native space of each functional run (using the inverse BOLD-to-T1w transformation). Components are also calculated separately within the WM and CSF masks. For each CompCor decomposition, the k components with the largest singular values are retained, such that the retained components’ time series are sufficient to explain 50 percent of variance across the nuisance mask (CSF, WM, combined, or temporal). The remaining components are dropped from consideration. The head-motion estimates calculated in the correction step were also placed within the corresponding confounds file. The confound time series derived from head motion estimates and global signals were expanded with the inclusion of temporal derivatives and quadratic terms for each (Satterthwaite et al., 2013). Frames that exceeded a threshold of 0.5 mm FD or 1.5 standardised DVARS were annotated as motion outliers. All resamplings can be performed with a single interpolation step by composing all the pertinent transformations (i.e. head-motion transform matrices, susceptibility distortion correction when available, and co-registrations to anatomical and output spaces). Gridded (volumetric) resamplings were performed using antsApplyTransforms (ANTs), configured with Lanczos interpolation to minimize the smoothing effects of other kernels (Lanczos, 1964). Non-gridded (surface) resamplings were performed using mri_vol2surf (FreeSurfer).

Many internal operations of fMRIPrep use Nilearn 0.6.1 (Abraham et al., 2014; RRID:SCR_001362), mostly within the functional processing workflow. For more details of the pipeline, see the section corresponding to workflows in fMRIPrep’s documentation.

### Regions of interest

We performed all analyses using an *a priori* region of interest (ROI) approach (Figure 2A). Using Freesurfer’s automated cortical and subcortical segmentation (Fischl, 2012), we targeted each participant’s hippocampus (HPC) and ventral temporal cortex (VTC; including bilateral parahippocampal gyrus, fusiform gyrus and inferior temporal gyrus; Ye et al., 2020). We also identified each participant’s retrosplenial cortex (RSC) and medial prefrontal cortex (mPFC), which were converted from the Schaefer atlas (Schaefer et al., 2018). All ROI masks were resampled to align with the resolution of the functional images.

To ensure the signal quality within the hippocampus, we removed individual voxels from each participant’s HPC mask that were extremely inactive and/or showed excessive temporal variability during encoding (Tambini & Davachi, 2013). Specifically, we excluded voxels if (1) their t statistics from trial-level Least Squares Separate modeling (see “*Least Squares Separate (LSS) modeling of encoding trials*”) was 2 standard deviation (SD) below the mean of the t statistics of all HPC voxels and/or (2) their temporal SD across all encoding trials was greater than 5SDs above the mean temporal SD across all HPC voxels (Tambini & Davachi, 2013). Additionally, we created anterior and posterior HPC masks by segmenting the whole HPC (after voxel exclusion) along the long axis and dividing the coronal slices of HPC into three sections, taking the most anterior third segment as the anterior HPC mask, and the most posterior third as posterior HPC.

### Statistical analyses

We performed statistical analyses using a combination of two-tailed paired t-tests, analysis of variance (ANOVA), and mixed-effects linear models. We adjusted p values and 95% confidence intervals with Bonferroni correction in all main analyses.

#### Behavioral performance

We computed each participant’s proportion correct (mean accuracy) and median response time (RT; correct trials only) on the associative recognition test for strong and weak memory conditions. We first compared the mean accuracy in each condition to the chance level (0.25) using a one-sample t-test, and then separately compared accuracy and RT across conditions using paired t-tests.

We also performed supplemental analyses to examine how accuracy and RT in strong and weak memory conditions might differ across the three visual categories. Specifically, we conducted 3×2 two-way ANOVAs with categories (objects vs. faces vs. scene) and encoding conditions (strong vs. weak) as factors.

#### Least Squares Separate (LSS) modeling of encoding trials

To capture replay of the encoded information during post-encoding rest periods, we first measured the multivoxel activity pattern associated with each encoding trial. Specifically, we used the LSS modeling approach and conducted a separate general linear model (GLM) for each encoding trial (Mumford et al., 2012, 2014), implemented using FEAT (Woolrich et al., 2001). Each model included a single encoding trial as the regressor of interest, and 4 regressors of no interest grouping all other trials by condition within the given encoding block (trials of the once present pairs; trials of the first, second, or third presentation of the thrice presented pairs). All trials were modeled as 4s boxcars convolved with hemodynamic response function (HRF). Each GLM also included six motion parameters, their 1^st^ temporal derivatives, and the top 5 anatomical CompCor separately from WM and CSF voxels (see “fMRI data preprocessing”) as nuisance regressors. In this way, we obtained voxelwise parameter estimates of activation that were specific to each encoding trial.

#### Replay analysis

We adapted the analyses from recent work to assess replay frequency of the encoding trials during post-encoding rest periods (Staresina et al., 2013; Schapiro et al., 2018). First, we further cleaned the signals in the preprocessed post-encoding rest scans by removing confounds including six motion parameters, their 1^st^ temporal derivatives, and the top 5 anatomical CompCor separately from WM and CSF voxels (see “fMRI data preprocessing”). For each participant and within each of the ROIs, we then computed pattern similarity scores (i.e., Pearson correlations) between the activation pattern of each trial in an encoding block and signals of each TR during the corresponding post-encoding rest period, yielding a matrix of pattern similarity scores for 72 encoding trials by 210 rest TRs for each pair of encoding-rest scans (Figure 2B&2C). Next, for each resulting matrix, we applied a threshold to each of the rest TRs, such that only the similarity scores that were greater than 1.5 SDs above the mean across all encoding trials within any given rest TR were considered as replay evidence of specific encoding trials (Figure 2C). Note that this SD thresholding approach was used as our primary method of identifying replay, and we focused on reporting results using this approach in the main text. However, in order to further validate our results, we also assessed replay with a supplemental thresholding approach and replicated our results (see “*Replay analysis with baseline rest thresholding approach*”).

After identifying the replay events during post-encoding rest, we then compared the replay frequency across strong and weak conditions within each ROI. Here, we first divided the encoding trials in the strong memory condition into three more fine-grained conditions according to the iteration of presentations (strong 1^st^ presentation, strong 2^nd^ presentation, strong last presentation). Each of the three fine-grained strong memory conditions contained the same number of trials as in the weak memory condition (18 trials). We then counted the number of TRs in each post-encoding rest period that showed replay for any encoding trial within each condition, and averaged the replay frequency across the two post-encoding rest periods. We then used a paired t-test to compare replay frequency between the strong memory (last presentation) condition and the weak memory condition, which was our main comparison of interest (see main text for reasoning). We also performed supplemental analyses separately examining for each ROI whether the difference in post-encoding replay between strong and weak memory conditions might depend on visual categories using two-way ANOVAs.

Importantly, we also compared the amount of replay across the three presentations in the strong memory condition in each ROI (averaged across two rests), using a one-way repeated-measures ANOVA. This analysis was conducted in order to test the hypothesis that the representations of the last presentation of the repeated pairs would be more replayed than the earlier presentations,

#### Replay analysis with baseline rest thresholding approach

To further validate our results obtained from the aforementioned SD thresholding approach (see “replay analysis”), we measured post-encoding replay with another thresholding approach adapted from prior work (Staresina et al., 2013). Here, we obtained a correlation matrix between each encoding block and the baseline rest scan for each subject and for each ROI. We then normalized the distribution of the pattern similarity scores of each encoding trial across all baseline rest TRs. This allowed us to create a reference distribution of ‘sham’ replay values for each encoding trial, as we assumed no replay should be observed in a baseline rest period prior to any encoding. We then took the 95% percentile point of the reference distribution as the replay threshold of a given encoding trial and applied these trial-specific thresholds to the correlation matrices relating encoding and post-encoding rest scans. Therefore, for each encoding trial, only TRs in the post-encoding rest period that showed a pattern similarity score greater than the trial-specific threshold value (corresponding to a p-value < 0.05 in the sham replay distribution) were considered as replay of that encoding trial. Separately for each ROI, we measured and compared post-encoding replay frequency (i.e., number of TRs that showed replay for at least one trial within a condition) obtained from this thresholding approach across the strong (last presentation) and weak memory conditions using a paired-sample t-test.

#### Controlling for encoding recency of strong and weak memory

To exclude the possibility that the prioritization of strong over weak memory during replay in the cortical regions was merely driven by encoding recency of the strong memory (last presentation) trials, we conducted a supplemental control analysis. This analysis compared replay frequency for a subset of trials across conditions that were matched in terms of the temporal distance between the encoding time and the onset of the rest period. To do this, we performed a median split of encoding time for trials in the weak memory condition for each participant and selected the half (9 trials) that were learned later, or more recently, during encoding. For each of these selected weak memory trials, we selected a strong memory last presentation trial within the same encoding block that was presented closest in time to, and *no later than*, the given weak memory trial. In this way, the selected strong memory trials were encoded approximately around the same time, but overall more distant to the rest period, as compared to the selected weak memory trials. This analysis included data from 34 encoding blocks in 25 participants (out of a total of 58 encoding blocks in 29 participants), as we were not able to get subsets of trials that fulfilled the search criteria in all participants’ data. We then examined and compared replay frequency across conditions using a paired t test separately for each cortical ROI.

#### Coordinated replay between the hippocampus and the cortex

To examine coordinated hippocampal-cortical replay of strong and weak memories, we assessed the proportion of encoding trials in each condition that were simultaneously replayed across regions. Given the segregation of connectivity between different HPC subregions to the cortex (Insausti & Muñoz, 2001; Kahn et al., 2008; Aggleton, 2012; Adnan et al., 2016; Ritchey & Cooper, 2020), this analysis was separately conducted for the anterior and posterior portions of the hippocampus. For each TR during rest and for each memory condition, we first extracted the binary vector of replay patterns across the encoding trials (1=replayed, 0=not replayed) from the thresholded similarity matrix of each brain region (Figure 2C right). We then computed a Jaccard similarity index between the patterns of each pair of the hippocampal subregion and the cortical region of interest (e.g., anterior HPC and VTC). We averaged the Jaccard similarity indices across all rest TRs in each condition as the measure of coordinated replay. We compared the mean Jaccard similarity index across the strong (last presentation) and weak memory conditions using a paired t-test.

We also examined whether the magnitude of difference in coordinated hippocampal-cortical replay between conditions varied across the anterior and posterior hippocampus. To do this, we examined the interaction between hippocampal subregion (anterior vs. posterior) and memory condition (strong vs. weak) with a two-way ANOVA for coordinated replay between hippocampus and each of the cortical ROIs.

#### Relating replay and behavioral performance

We performed trial-level mixed-effects linear models to examine the association between post-encoding replay and subsequent memory outcomes on the recognition test. Here, we counted the number of TRs each encoding trial was replayed across the entire rest period. Note that for pairs in the strong memory condition, we only included trials within the strong memory last presentation condition. In each ROI, we predicted whether a pair was remembered or forgotten on the recognition test (i.e., retrieval success: correct vs. incorrect) with the replay frequency of the given trial during post-encoding rest (mean-centered), the encoding condition of the pair (strong vs. weak; we separately treated each condition as the baseline in parallel models with identical terms), the univariate activity associated with the encoding trial (mean-centered), as well as their interactions. Univariate encoding activity for each trial was measured by computing the average of the trial-specific parameter estimates obtained from LSS modeling across all voxels within each ROI. All fixed effects (including the intercept) were included as random effects, grouped by participant. Models used a logistic linking function.

Following up on the finding that replay in the whole HPC and anterior HPC was significantly associated with the memory outcomes of weakly encoded information, we performed an exploratory analysis asking whether this association might be stronger in people with overall poor versus good memory performance. We first performed a median split of recognition accuracy in the weak memory condition across participants (Figure S4, left). Then within each of whole HPC and anterior HPC, we performed a trial-level mixed-effects linear model predicting the memory outcome of each pair in the weak memory condition (correct vs. incorrect) with the replay frequency of that pair (mean-centered), the participant’s performance level (good vs. poor; we separately treated each level as the baseline in parallel models with identical terms) and their interaction. Replay frequency and the intercept were included as random effects in the models, grouped by participant. Models used a logistic linking function.

## Acknowledgements

We thank current and former members of the Davachi Lab for their constructive feedback. This work is supported by the National Institutes of Health grant R01-MH0746492.

