## SupplementalInfo for "Repetition accelerates neural markers of memory consolidation"

### Supplemental Information

#### Supplemental Data

Behavioral performance

We examined whether memory performance on the associative recognition test varied across encoding conditions (within-subjects factor) and visual categories (across-subjects factor) using a two-way ANOVA. Results revealed a significant main effect of encoding condition ($F(1,52)=21.70$, $p<0.001$, $\eta^2=0.29$), such that performance was significantly higher in the strong memory condition than the weak memory condition. No significant main or interactive effect of category was found (both $p>0.13$).

Within the correct trials, participants had a median RT of 1.50 s (SD=0.55) in the strong memory condition, which was significantly faster than in the weak memory condition (Mean=1.77 s, SD=0.52; $t(28) = -4.46$, $p<0.001$, 95% CI[-0.38, -0.14], Cohen's $d=0.83$). A two-way ANOVA comparing RT across encoding conditions and visual categories revealed a marginal main effect of encoding condition ($p=0.07$) while neither the main nor interactive effect of category was significant (both $p>0.33$).

#### Supplemental Tables

**Table S1-1, Related to Figure 5. Predicting memory outcomes with replay frequency in the hippocampus.**
We performed a multilevel mixed-effects linear model to assess the association between trial-specific replay frequency in the hippocampus and trial-to-trial retrieval success on the associative recognition test, while controlling for the univariate encoding activity associated with the trial. Parallel models with identical terms were run separately treating weak memory condition (Table S1-1) or strong memory condition as baseline (Table S1-2). Additional models were run predicting retrieval success separately with anterior or posterior hippocampal replay frequency, and all other terms in these models were identical to the whole hippocampus model. ~$p<0.1$, *$p<0.05$, **$p<0.01$, ***$p < 0.001$ (statistical significance and 95% confidence interval adjusted with Bonferroni correction).

*retrieval success~replay frequency*encoding univariate*encoding condition + (replay frequency*encoding univariate* encoding condition|subject)*

*Whole hippocampus*

| | **HPC (baseline: weak memory condition)** | | |
|---|---|---|---|
| *Predictors* | *Estimate* | *CI* | *p* |
| (Intercept) | 1.01085 | 0.62169 – 1.40001 | <0.001*** |
| Replay frequency | 0.04372 | 0.00277 – 0.08468 | 0.036* |
| Encoding univariate | 0.00465 | -0.00443 – 0.01373 | 0.315 |
| Encoding condition | 1.81319 | 1.26977 – 2.35661 | <0.001*** |
| Replay frequency*Encoding univariate | -0.00006 | -0.00136 – 0.00124 | 0.929 |
| Replay frequency*Encoding condition | -0.01321 | -0.07996 – 0.05354 | 0.698 |
| Encoding univariate*Encoding condition | -0.00969 | -0.02540 – 0.00602 | 0.227 |
| Replay frequency*Encoding univariate* Encoding condition | 0.00030 | -0.00178 – 0.00238 | 0.778 |
| $N_{sub}$ | 29 | | |
| Observations | 1044 | | |

*Hippocampal subregions*

| | Anterior HPC (baseline: weak memory condition) | | |
|---|---|---|---|
| *Predictors* | *Estimate* | *CI* | *p* |
| (Intercept) | 0.98847 | 0.54112 – 1.43582 | <0.001*** |
| Replay frequency | 0.05601 | 0.01519 – 0.09683 | 0.004** |
| Encoding univariate | 0.00361 | -0.00289 – 0.01011 | 0.427 |
| Encoding condition | 1.78178 | 1.16351 – 2.40005 | <0.001*** |
| Replay frequency*Encoding univariate | 0.00033 | -0.00054 – 0.00119 | 0.794 |
| Replay frequency*Encoding condition | -0.06114 | -0.13038 – 0.00811 | 0.096~ |
| Encoding univariate*Encoding condition | -0.01072 | -0.02354 – 0.00210 | 0.121 |
| Replay frequency*Encoding univariate* Encoding condition | 0.00046 | -0.00107 – 0.00198 | 1 |

| | Posterior HPC (baseline: weak memory condition) | | |
|---|---|---|---|
| *Predictors* | *Estimate* | *CI* | *p* |
| (Intercept) | 0.96350 | 0.52259 – 1.40440 | <0.001*** |
| Replay frequency | -0.00795 | -0.06804 – 0.05214 | 1 |
| Encoding univariate | -0.00248 | -0.01100 – 0.00605 | 1 |
| Encoding condition | 1.82789 | 1.22485 – 2.43093 | <0.001*** |
| Replay frequency*Encoding univariate | -0.00039 | -0.00241 – 0.00162 | 1 |
| Replay frequency*Encoding condition | 0.08597 | -0.01940 – 0.19134 | 0.135 |
| Encoding univariate*Encoding condition | -0.00555 | -0.02383 – 0.01272 | 0.991 |
| Replay frequency*Encoding univariate* Encoding condition | 0.00112 | -0.00302 – 0.00527 | 1 |
| $N_{sub}$ | 29 | | |
| Observations | 1044 | | |

**Table S1-2, Related to Figure 5. Outputs of models parallel to Table S1-1, with strong memory condition treated as baseline.**

*Whole hippocampus*

| | **HPC (baseline: strong memory condition)** | | |
|---|---|---|---|
| *Predictors* | *Estimate* | *CI* | *p* |
| (Intercept) | 2.54582 | 2.07062 – 3.02103 | <0.001*** |
| Replay frequency | 0.02247 | -0.03389 – 0.07882 | 0.435 |
| Encoding univariate | -0.00567 | -0.01785 – 0.00651 | 0.361 |
| Encoding condition | -1.45481 | -1.89558 – -1.01404 | <0.001*** |
| Replay frequency*Encoding univariate | 0.00067 | -0.00109 – 0.00242 | 0.456 |
| Replay frequency*Encoding condition | 0.03059 | -0.03596 – 0.09715 | 0.368 |
| Encoding univariate*Encoding condition | 0.01129 | -0.00419 – 0.02677 | 0.153 |
| Replay frequency*Encoding univariate* Encoding condition | -0.00099 | -0.00343 – 0.00146 | 0.429 |
| $N_{sub}$ | 29 | | |
| Observations | 1044 | | |

*Hippocampal subregions*

| | **Anterior HPC (baseline: strong memory condition)** | | |
|---|---|---|---|
| *Predictors* | *Estimate* | *CI* | *p* |
| (Intercept) | 2.53089 | 1.98905 – 3.07274 | <0.001*** |
| Replay frequency | -0.00295 | -0.05887 – 0.05296 | 1 |
| Encoding univariate | -0.00674 | -0.01732 – 0.00384 | 0.307 |
| Encoding condition | -1.47150 | -1.99055 – -0.95246 | <0.001*** |

| | | | |
|---|---|---|---|
| Replay frequency*Encoding univariate | 0.00084 | -0.00048 – 0.00216 | 0.311 |
| Replay frequency*Encoding condition | 0.07042 | 0.00038 – 0.14045 | 0.048* |
| Encoding univariate*Encoding condition | 0.01060 | -0.00197 – 0.02317 | 0.118 |
| Replay frequency*Encoding univariate* Encoding condition | -0.00071 | -0.00247 – 0.00106 | 0.737 |

| | **Posterior HPC (baseline: strong memory condition)** | | |
|---|---|---|---|
| *Predictors* | *Estimate* | *CI* | *p* |
| (Intercept) | 2.56240 | 2.01580 – 3.10899 | <0.001*** |
| Replay frequency | 0.07481 | -0.01753 – 0.16715 | 0.139 |
| Encoding univariate | -0.00774 | -0.02314 – 0.00767 | 0.520 |
| Encoding condition | -1.55319 | -2.06269 – -1.04370 | <0.001*** |
| Replay frequency*Encoding univariate | 0.00048 | -0.00293 – 0.00389 | 1 |
| Replay frequency*Encoding condition | -0.07917 | -0.18092 – 0.02259 | 0.162 |
| Encoding univariate*Encoding condition | 0.00554 | -0.01226 – 0.02334 | 0.970 |
| Replay frequency*Encoding univariate* Encoding condition | -0.00105 | -0.00522 – 0.00313 | 1 |
| $N_{sub}$ | 29 | | |
| Observations | 1044 | | |

**Table S2-1, Related to Figure 5. Predicting memory outcomes with replay frequency in cortical regions.**

We performed a multilevel mixed-effects linear model predicting trial-to-trial retrieval success on the memory test with trial-specific replay frequency in each cortical region of interest. All other model setup was identical to the aforementioned model with hippocampal replay. Table S2-1 shows outputs of models treating weak memory condition as the baseline condition, and Table S2-2 shows outputs of models treating strong memory condition as the baseline condition. ~p<0.1, *p<0.05, **p<0.01, ***p < 0.001 (statistical significance and 95% confidence interval adjusted with Bonferroni correction).

*retrieval success~replay frequency*encoding univariate*memory condition + (replay frequency*encoding univariate*memory condition|subject)*

| | **VTC (baseline: weak memory condition)** | | |
|---|---|---|---|
| *Predictors* | *Estimate* | *CI* | *p* |
| (Intercept) | 0.94680 | 0.47223 – 1.42136 | <0.001*** |
| Replay frequency | 0.02067 | -0.00594 – 0.04729 | 0.189 |
| Encoding univariate | 0.00483 | -0.00367 – 0.01332 | 0.521 |
| Encoding condition | 1.77840 | 1.12731 – 2.42949 | <0.001*** |
| Replay frequency*Encoding univariate | -0.00006 | -0.00059 – 0.00046 | 1 |
| Replay frequency*Encoding condition | -0.02509 | -0.06985 – 0.01968 | 0.539 |
| Encoding univariate*Encoding condition | -0.01238 | -0.02687 – 0.00210 | 0.122 |
| Replay frequency*Encoding univariate* Memory condition | -0.00017 | -0.00122 – 0.00087 | 1 |

| | **RSC (baseline: weak memory condition)** | | |
|---|---|---|---|
| *Predictors* | *Estimate* | *CI* | *p* |
| (Intercept) | 0.99149 | 0.51462 – 1.46836 | <0.001*** |
| Replay frequency | -0.00856 | -0.03608 – 0.01897 | 1 |
| Encoding univariate | 0.00238 | -0.00471 – 0.00946 | 1 |

| | | | |
|---|---|---|---|
| Encoding condition | 1.73630 | 1.09460 – 2.37799 | <0.001*** |
| Replay frequency*Encoding univariate | -0.00065 | -0.00117 – -0.00012 | 0.009** |
| Replay frequency*Encoding condition | 0.03225 | -0.02128 – 0.08578 | 0.448 |
| Encoding univariate*Encoding condition | -0.00284 | -0.01518 – 0.00951 | 1 |
| Replay frequency*Encoding univariate* Encoding condition | 0.00101 | -0.00010 – 0.00212 | 0.089~ |

| | **mPFC (baseline: weak memory condition)** | | |
|---|---|---|---|
| *Predictors* | *Estimate* | *CI* | *p* |
| (Intercept) | 0.97555 | 0.50748 – 1.44362 | <0.001*** |
| Replay frequency | 0.00759 | -0.01819 – 0.03336 | 1 |
| Encoding univariate | 0.00038 | -0.00430 – 0.00505 | 1 |
| Encoding condition | 1.85556 | 1.19519 – 2.51593 | <0.001*** |
| Replay frequency*Encoding univariate | -0.00026 | -0.00071 – 0.00020 | 0.537 |
| Replay frequency*Encoding condition | 0.02544 | -0.02608 – 0.07696 | 0.712 |
| Encoding univariate*Encoding condition | -0.00504 | -0.01700 – 0.00693 | 0.941 |
| Replay frequency*Encoding univariate* Encoding condition | 0.00013 | -0.00082 – 0.00108 | 1 |
| $N_{sub}$ | 29 | | |
| Observations | 1044 | | |

**Table S2-2, Related to Figure 5. Outputs of models parallel to Table S2-1, with strong memory condition treated as baseline.**

| | **VTC (baseline: strong memory condition)** | | |
|---|---|---|---|
| *Predictors* | *Estimate* | *CI* | *p* |
| (Intercept) | 2.46411 | 1.89197 – 3.03625 | <0.001*** |

| | | | |
|---|---|---|---|
| Replay frequency | -0.00459 | -0.03895 – 0.02977 | 1 |
| Encoding univariate | -0.00701 | -0.01828 – 0.00426 | 0.410 |
| Encoding condition | -1.45870 | -2.00467 – -0.91273 | <0.001*** |
| Replay frequency*Encoding univariate | -0.00022 | -0.00109 – 0.00065 | 1 |
| Replay frequency*Encoding condition | 0.02737 | -0.02000 – 0.07474 | 0.500 |
| Encoding univariate*Encoding condition | 0.01228 | -0.00203 – 0.02659 | 0.120 |
| Replay frequency*Encoding univariate* Encoding condition | 0.00011 | -0.00093 – 0.00115 | 1 |

| | **RSC (baseline: strong memory condition)** | | |
|---|---|---|---|
| *Predictors* | *Estimate* | *CI* | *p* |
| (Intercept) | 2.49162 | 1.91406 – 3.06919 | <0.001*** |
| Replay frequency | 0.02204 | -0.02205 – 0.06614 | 0.694 |
| Encoding univariate | -0.00161 | -0.01122 – 0.00801 | 1 |
| Encoding condition | -1.45894 | -1.99868 – -0.91920 | <0.001*** |
| Replay frequency*Encoding univariate | 0.00041 | -0.00056 – 0.00139 | 0.926 |
| Replay frequency*Encoding condition | -0.03138 | -0.08403 – 0.02127 | 0.461 |
| Encoding univariate*Encoding condition | 0.00378 | -0.00836 – 0.01591 | 1 |
| Replay frequency*Encoding univariate* Encoding condition | -0.00109 | -0.00219 – 0.00001 | 0.054~ |

| | **mPFC (baseline: strong memory condition)** | | |
|---|---|---|---|
| *Predictors* | *Estimate* | *CI* | *p* |
| (Intercept) | 2.54722 | 1.96372 – 3.13072 | <0.001*** |
| Replay frequency | 0.02784 | -0.01348 – 0.06917 | 0.320 |
| Encoding univariate | -0.00445 | -0.01291 – 0.00401 | 0.623 |

| | | | |
|---|---|---|---|
| Encoding condition | -1.53352 | -2.08803 – -0.97901 | <0.001*** |
| Replay frequency*Encoding univariate | -0.00032 | -0.00107 – 0.00043 | 0.908 |
| Replay frequency*Encoding condition | -0.01997 | -0.06933 – 0.02938 | 0.998 |
| Encoding univariate*Encoding condition | 0.00484 | -0.00493 – 0.01461 | 0.707 |
| Replay frequency*Encoding univariate* Encoding condition | 0.00005 | -0.00084 – 0.00093 | 1 |
| $N_{sub}$ | 29 | | |
| Observations | 1044 | | |

**Table S3-1, related to Figure S4. Predicting weak memory outcomes with trial-specific hippocampal replay frequency across good and poor performers.**

We performed an exploratory analysis using multilevel mixed-effects linear models to examine whether the significant association between retrieval success for the weakly encoded pairs and trial-specific hippocampal replay frequency differs across good and poor performers. Parallel models with identical terms were run separately treating either the poor (Table S3-1) or good (Table S3-2) performer group as the baseline. Additional models were performed with replay frequency in the anterior HPC, given that replay frequency in the anterior portion of the hippocampus was significantly associated with weak memory outcomes in previous results (see Table S1-1). ~p<0.1; *p<0.05; **p<0.01; ***p < 0.001.

*retrieval success~replay frequency* memory performance+ (replay frequency |subject)*

| | **HPC (baseline: poor performer group)** | | |
|---|---|---|---|
| *Predictors* | *Estimate* | *CI* | *p* |
| (Intercept) | 0.11 | -0.16 – 0.37 | 0.432 |
| Replay frequency | 0.06 | 0.01 – 0.11 | 0.020* |
| Memory performance | 1.68 | 1.24 – 2.12 | <0.001*** |
| Replay frequency*Memory performance | -0.07 | -0.14 – 0.01 | 0.072~ |
| | **Anterior HPC (baseline: poor performer group)** | | |
| *Predictors* | *Estimate* | *CI* | *p* |
| (Intercept) | 0.08 | -0.18 – 0.34 | 0.546 |
| Replay frequency | 0.06 | 0.01 – 0.11 | 0.010* |
| Memory performance | 1.71 | 1.26 – 2.15 | <0.001*** |
| Replay frequency*Memory performance | -0.04 | -0.11 – 0.04 | 0.324 |
| $N_{sub}$ | 29 | | |
| Observations | 522 | | |

**Table S3-2, related to Figure S4. Outputs of models parallel to Table S3-1, with strong memory condition treated as baseline.**

| | **HPC (baseline: good performer group)** | | |
|---|---|---|---|
| *Predictors* | *Estimate* | *CI* | *p* |
| (Intercept) | 1.79 | 1.43 – 2.15 | <0.001*** |
| Replay frequency | -0.01 | -0.07 – 0.05 | 0.719 |
| Memory performance | -1.68 | -2.12 – -1.24 | <0.001*** |
| Replay frequency*Memory performance | 0.07 | -0.01 – 0.14 | 0.072~ |
| | **Anterior HPC (baseline: good performer group)** | | |
| *Predictors* | *Estimate* | *CI* | *p* |
| (Intercept) | 1.79 | 1.43 – 2.15 | <0.001*** |
| Replay frequency | 0.02 | -0.03 – 0.08 | 0.417 |
| Memory performance | -1.71 | -2.15 – -1.26 | <0.001*** |
| Replay frequency*Memory performance | 0.04 | -0.04 – 0.11 | 0.324 |
| $N_{sub}$ | 29 | | |
| Observations | 522 | | |

#### Supplemental Figures

##### Figure S1, Related to Figure 3. Post-encoding replay: encoding condition × image category

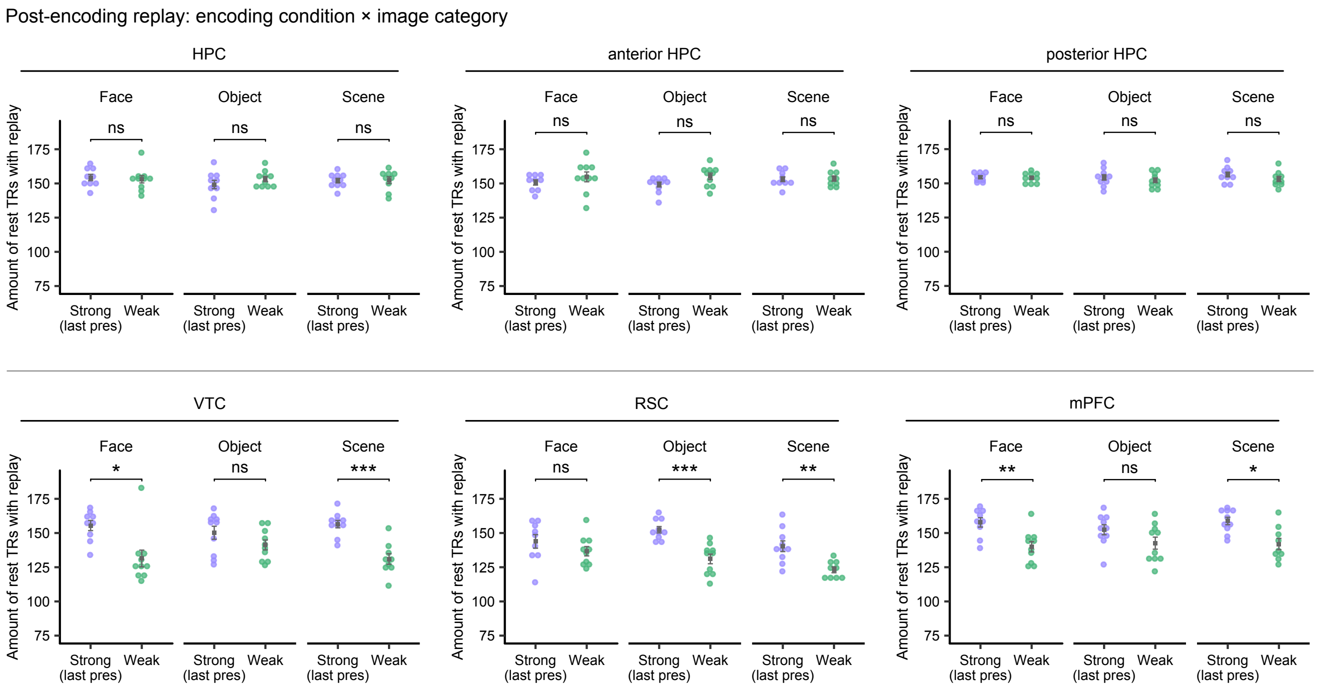


A two-way ANOVA comparing replay frequency across encoding conditions (within-subjects factor) and visual categories (across-subjects factor) was conducted in each ROI. Results revealed no significant main or interactive effect of either variable in HPC (all $p>0.53$) or posterior HPC (all $p>0.15$). Anterior HPC showed a trend towards a significant main effect of encoding condition ($F(1,52)=3.46$, $p=0.068$, $\eta^2=0.062$; the follow-up two-sample t-test within each visual category did not reveal any significant difference across strong and weak memory conditions, all $p>0.17$).

All cortical ROIs showed a significant main effect of encoding condition (VTC: $F(1,52)= 30.04$, $p<0.001$, $\eta^2=0.37$; RSC: $F(1,52)= 28.99$, $p<0.001$, $\eta^2=0.36$; mPFC: $F(1,52)= 23.45$, $p<0.001$, $\eta^2=0.31$). For replay in VTC and mPFC, the comparison between strong and weak memories within each visual category revealed a significant difference in face and scene categories (all $p<0.014$), but not in the object category ($p=0.47$). For replay in RSC, the difference between strong and weak memories was significant in object and scene categories (both $p<0.0053$), but not the face category ($p=0.76$).

A significant main effect of visual category was only found in RSC ($F(2,52)= 4.79$, $p=0.01$, $\eta^2=0.16$, pairwise comparisons did not reveal a significant difference between any two categories). No other regions showed a significant main or interactive effect of visual category (all $p>0.10$).

Each dot represents a participant. Squared dots indicate mean values. Error bars indicate standard error. ns: not significant; **$p<0.01$; ***$p < 0.001$ (statistical significance of all follow-up t-tests adjusted with Bonferroni correction).

**Figure S2, Related to Figure 3. Results of post-encoding replay with baseline rest thresholding approach**

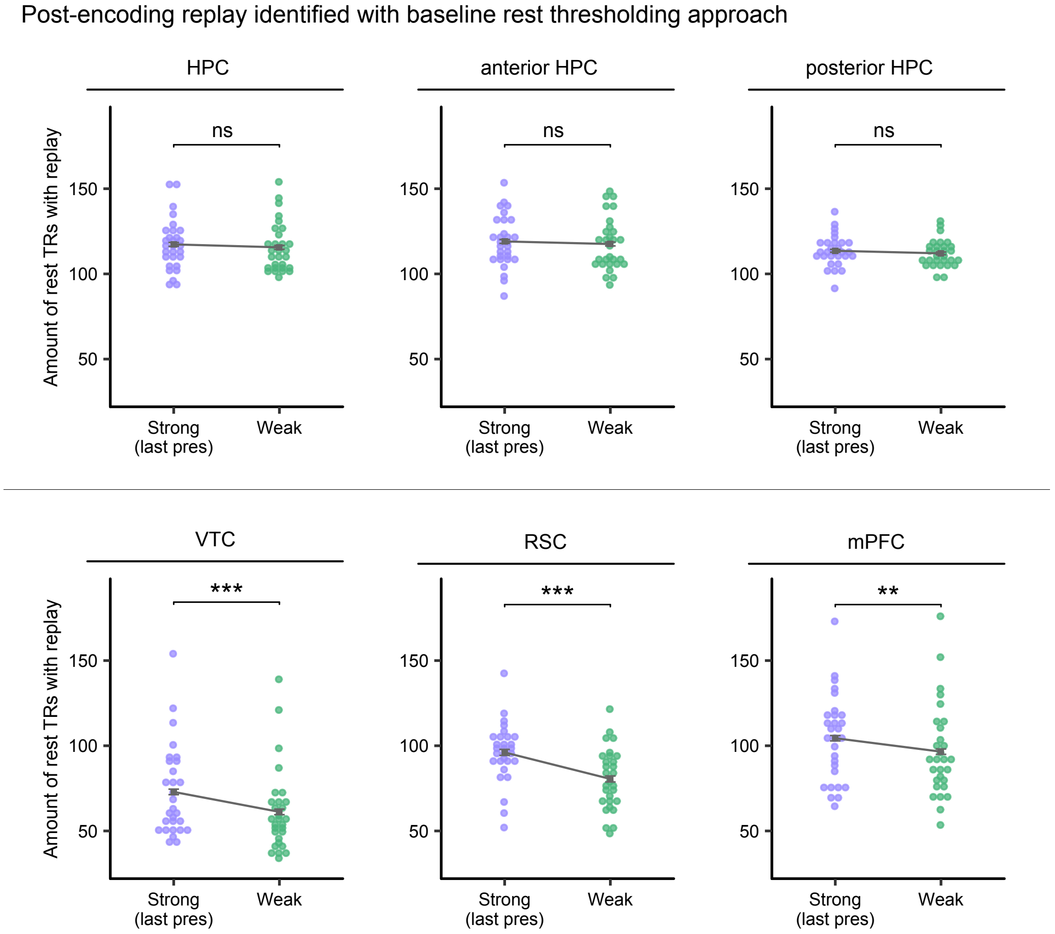


Supplementary analysis comparing post-encoding replay across strong and weak memory conditions with replay identified using the baseline rest thresholding approach. We found no differential replay for strong and weak memories in HPC (t (28)=1.08, p=0.29, 95% CI[-1.56, 5.01]), anterior HPC (t (28)=0.95, p=0.35, 95% CI[-1.74, 4.78]), or posterior HPC (t (28)=1.28, p=0.21, 95% CI[-0.91, 3.94]). Significantly higher replay frequency for strong versus weak memory was found in all cortical ROIs (VTC: t (28)=5.10, p<0.001, 95% CI[7.01, 16.44], Cohen's d=0.95; RSC: t (28)=6.26, p<0.001, 95% CI[10.49, 20.69], Cohen's d=1.16; mPFC: t(28)=3.54, p=0.001, 95% CI[3.33, 12.46], Cohen's d=0.66). Each dot represents a participant, squared dots indicate mean values. Error bars show within-subject standard error. ns: not significant; **p<0.01; ***p < 0.001.

**Figure S3, Related to Figure 3. Cortical replay of strong and weak memories matched in encoding recency**

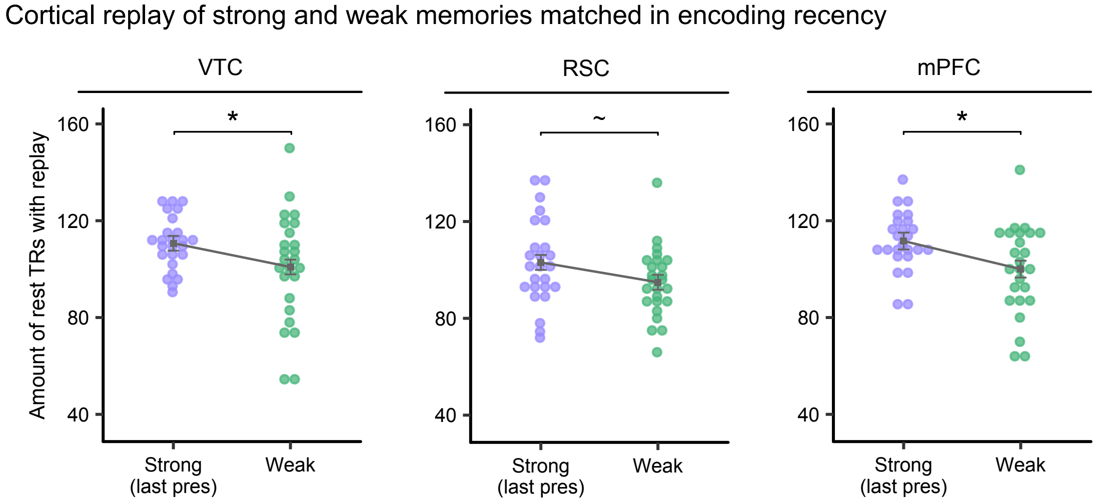


Supplementary control analysis examining cortical replay of subsets of strong and weak memory trials matched in encoding recency. The patterns of replay across strong (last presentation) and weak memory conditions persisted in cortical ROIs after controlling for encoding recency (VTC: t (24)=2.27, p=0.03, 95% CI[0.90, 18.70], Cohen's d=0.45; RSC: t (24)=1.87, p=0.07, 95% CI[-0.84,17.24], Cohen's d=0.37; mPFC: t (24)=2.35, p=0.03, 95% CI[1.43, 21.77], Cohen's d=0.47). Each dot represents a participant, squared dots indicate mean values. Error bars show within-subject standard error. ~p<0.1; *p<0.05.

**Figure S4, Related to Figure 5. Predicting weak memory outcome with post-encoding replay in good vs. poor performers.**

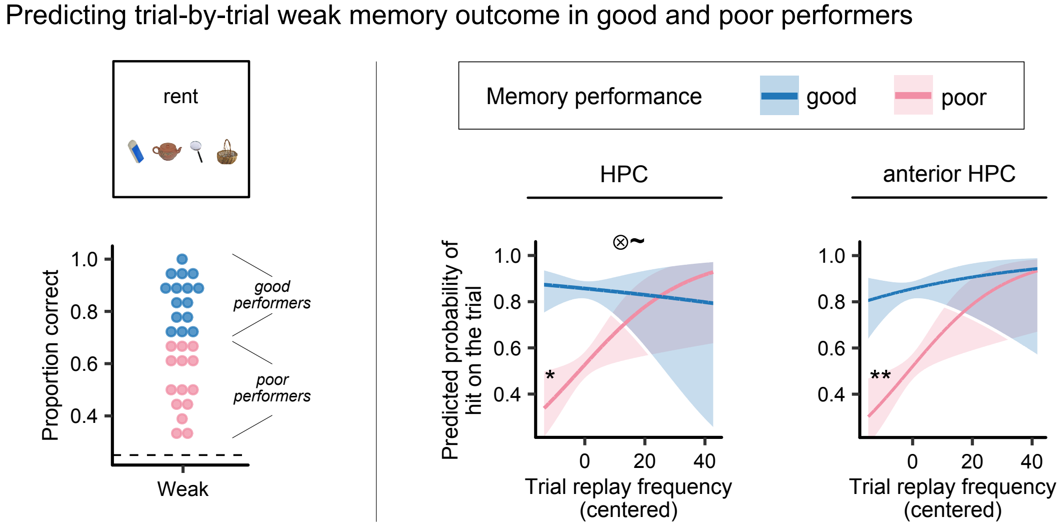


Exploratory analysis predicting memory outcome of each pair in the weak memory condition with the pair-specific hippocampal replay frequency and memory performance group. Performance group was defined according to the median split of associative recognition accuracy in the weak memory condition. We found that for poor performers, higher replay frequency of a weakly encoded pair during rest was significantly associated with greater probability of remembering that pair on the memory test (HPC: b=0.057, p=0.020; anterior HPC: b=0.062, p=0.0097; Table S3-1), while this association was not found in good performers (all p>0.41; Table S3-2). Further, the association between trial-specific hippocampal replay frequency and memory outcome was also marginally strong for poor performers than good performers (b=-0.068, p=0.071; S3-1). See Table S3-1&3-2 for full model outputs. Ribbons represent 95% confidence intervals. ⊗: interaction; ~p<0.1; *p<0.05; **p<0.01.
